# Mechanical Stress dissipation in locally folded epithelia is orchestrated by calcium waves and nuclear tension changes

**DOI:** 10.1101/2025.04.04.647200

**Authors:** Marvin Brun-Cosme-Bruny, Lydia Pernet, Kenny Elias, Christophe Guilluy, Christiane Oddou, Monika Elzbieta Dolega

## Abstract

Epithelia are continuously exposed to a range of biomechanical forces such as compression, stretch and shear stress arising from their dynamic microenvironments and associated to their function. Changes in tension such as stretch are known to trigger cell rearrangements and divisions, and impact cellular transcription until mechanical stress is dissipated. How cells process, adapt and respond to mechanical stress is being intensively investigated. In here we focus on epithelial folding which is the fundamental process of transformation of flatmonolayers into 3D functional tissues. By combining the innovative method for fold generation, live imaging, mechanobiology tools and chemical screening, we uncover the role of calcium waves on mechanical adaptation of folded epithelia that occurs at the tissue and nuclear level. Folding associated tensional load results in the nuclear flattening which is recovered in the time scale of minutes and is dependent on the calcium wave that spread outwards from the channel and across the epithelium. By creating a mutant overexpressing LBR that relaxed nuclear envelope, we demonstrated that despite presence of calcium waves, nuclear tension increase was essential to trigger nuclear shape recovery post folding through the activation of cellular contractility in the cPLA2 dependent manner. Overall our results identify the molecular mechanism for nuclear shape recovery and indicate that mechanical stress dissipation program is activated at the level of nuclei which serve as internal tension sensors.

## Introduction

The formation of epithelial folds is a highly dynamic fundamental process in development that converts initially flat sheets of tightly interconnected cells into a 3D functional tissues and organs. *In vivo* models of embryonic development enabled our understanding of molecular pathways regulating folding and cellular patterning and in parallel underlined the importance of mechanical cues in the initiation and progression of morphogenesis processes. Indeed, folding is a physical process that requires mechanical force asymmetries to generate an invagination ^1^. Drosophila ventral furrow formation is one of the best characterized models of epithelial folding, where a thin strip of prospective furrow cells accumulates myosin on their apical side, which through the active process of apical constriction leads to the fold formation and mesoderm internalization ^2^. Mechanical forces driving furrow formation result in extreme cell shape changes that can elongate up to 1.7 times ^3^, which as well impose nuclear deformations and determine transcription programs ^4^. Whereas initiation of the process happens locally along the ventral axis, increasing evidence shows that a biophysical embryo-scale coordination is crucial to complete the gastrulation process and to ensure epithelial integrity under mechanically dynamic and challenging conditions^5^. However, precise physical and molecular mechanisms of intercellular communication and coordination remain still poorly studied.

Calcium is an ubiquitous and universal second messenger that impacts fundamental cellular processes such as transcription ^6^, cell cycle progression^7^, apoptosis ^8^, cytoskeleton redistribution and the generation of forces through the acto-myosin contractility^9^. Calcium signaling is mediated by calcium-sensing effectors that translate calcium signals with their characteristic spatio-temporal dynamics into a specific cellular response ^10–12^. A small stimulus will activate low number of calcium channels which will result in a transient and local increase, whereas high amplitude entry of calcium can activate further the calcium channels on endoplasmic reticulum, such as IP3R or RyR channel, and cause a general spike in calcium through a positive feedback loop. Calcium signals are known to be activated by mechanical forces through the mechano-sensitive channels such as Piezo or TRPV ^13^. The effective calcium concentration changes are transient, but can last from a single spike of miliseconds up to a global increase lasting over hours. This wide temporal range enables activating a whole subset of responses in the cell ^12^. From the perspective of intercellular communication calcium has the capacity to be rapidly transmitted between the cells interconnected through the gap-junctions. Indeed, calcium waves were observed in numerous developmental processes and perturbation of intracellular calcium concentrations has impeded the proper development such as during gastrulation of sea urchin ^14^, Xenopus ^15^, and zebrafish ^16^. Live imaging of early cleavage of zebrafish embryos showed that localized calcium transients accompany furrow positioning, and through induction, propagation and deepening of calcium waves ^17^. Moreover, calcium waves are equally important for maintenance of tissue integrity as they play a significant role in the actin-cable formation at the scar during the wound healing process ^18^. Altogether, these correlative observations underlie the fundamental role of calcium waves in numerous physiological processes in development and homeostasis, but the molecular understanding of the role they play in these processes is still missing mainly because of the systems complexity where cellular behaviors are determined by an interplay between mechanical forces and genetic programs.

Here, we employ a microsystem that enables recapitulating the characteristic shape changes of the cells and the global 3D curvature that is present during the process of *Drosophila* gastrulation ^19^. With our system, in which tissue mechanical variables (in-plane strain, curvature) are well characterized and controlled externally, we probe how local epithelial deformation is perceived and transmitted to the neighboring cells to collectively organize the tissue response. As in the system we start from the initially flat and homogenous monolayer, the experimental setup allows investigating the intercellular communication and collective behavior originating solely from the local alteration of the mechanical state. We showed previously that triggered folds are accompanied by the spreading of calcium waves outwards from the folds, yet their mechanism of initiation and role in epithelial collective response is missing. Here, by combining pharmacological treatments targeting calcium signaling and numerical modelling, we characterized the mechanisms of calcium wave initiation and intercellular propagation. We further observed that in the timescale of minutes nuclei being initially deformed upon folding partially recovered their shape but solely in the presence of calcium. Moreover, based on FLIM-Flipper-TR microscopy we showed that formed folds caused an increase in the nuclear envelope tension of the underlying cells. By creating a mutant overexpressing LaminB receptor (LBR) that relaxed nuclear envelope, we demonstrated that despite the presence of calcium waves, nuclear tension increase was essential to trigger nuclear shape recovery post folding through the activation of cellular contractility in the cPLA2 dependent manner. Overall, our results identify molecular mechanism for nuclear shape recovery and uncover the role of the calcium waves in the mechanical stress dissipation program, which as we showed is activated at the level of nuclei that serve as internal tissue tension sensors.

## RESULTS

### Local epithelial folding triggers calcium waves that are necessary for the deformation-associated nuclear shape recovery

To dissect mechanical cues from biochemical signaling, we used a microfluidic system that experimentally triggers epithelial folds within a homogeneous monolayer of epithelial cells to recapitulate stereotypic deformations observed *in vivo* (Figure 1A-C)^19^. The microsystem is composed of a single channel of 400 µm width enclosed with a 20µm thick PDMS membrane and an inlet (Figure 1A) through which a controlled vacuum is exerted to create concave membrane deformation. Epithelial monolayer growing on top of the membrane is deformed mechanically through the changes of the pressure at the inlet (Figure 1Aα). We measured that for the pressure of −450 mbar we recapitulate typical gradient of cell deformation in *Drosophila* gastrulation ^20^ which is present at the border of the channel where the curvature is the highest (Figure 1B). We observed that such strong deformations cause calcium increase in the cells above the channel, and the calcium wave propagated (Figure 1C) from the channel border outwards to non-deformed cell reaching a distance of about 200 µm (Figure 1D). Analysis of intensity of calcium marker Fluo4AM in time quantitatively confirmed the gradual propagation of calcium wave (Figure 1E).

**Figure 1.**
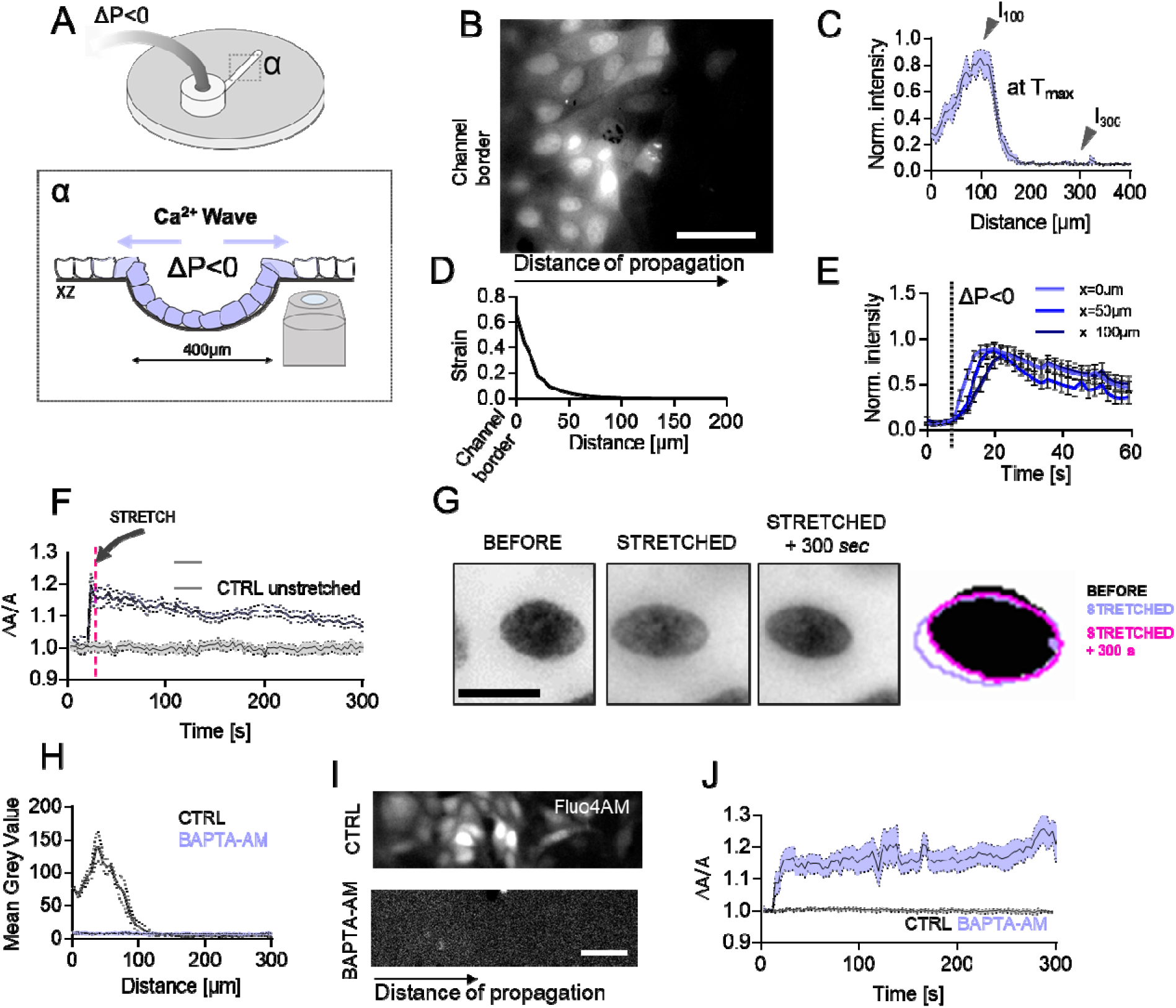
Local epithelial folding triggers calcium waves that are necessary for deformation-associated nuclear shape recovery. A) Schematic representation of the microfluidic system used to induce epithelial folds from a 2D epithelium. *α* shows the dimensions of the channel with the observation zone that is next to the channel border. Calcium waves are generated above the channel and spread laterally. B) Epifluorescence image from the live observation of spreading Ca2+ wave. Cells were stained with Ca^2+^ indicator Fluo4AM (image represents a time-point at t=18sec); Scale 50 µm. C) Quantification of calcium waves propagation distance based on the measurement of fluorescence intensity and normalized by the maximum value of Intensity at T_max_ (I_max_); Error bar: SEM, N=6 independent experiments D) Distribution of strain associated with the epithelial folding, x=0 for the channel border and x>0 for the epithelium. Image from B) corresponds to the same distance as on the X axis in D). E) Quantification of calcium signal intensity change as a function of time and with respect to the position with x=0 at channel border, and 50 and 100um from the channel. Error bar: SEM, N=6 independent experiments F) Quantification of the projected nuclear area change in time for cells that were stretched (next to the channel) and the cells at 800 µm away from the channel (unstreched). Results are normalized to A_0_, the nucleus area before the folds were induced. Error bar: SEM, N=9 independent experiments G) Microphotographs of inverted fluorescence of Hoechst stained nuclei before, just after epithelial fold was induced, and 300 s later. Color masks super-positioned represent the nuclear area measured before stretch, just after stretch and 300 s later; Scale 10 µm H) Quantification of the mean Grey value of Fluo4AM as a function of distance for control cells (as in C) and for cells pre-treated with BAPTA-AM. Error bar: SEM, N=6 independent experiments I) Images showing Fluo4AM staining of control and BAPTA-AM treated cells upon epithelial folding. Error bar: SEM, N=3 independent experiments; Scale 20 µm J) Quantification of the nuclear area change for control and BAPTA-AM treated cells upon folding; Error bar: SEM, N=3 independent experiments

Closer inspection of nuclei which were strongly stained by Fluo4AM revealed their initial elongation upon folding (increase in projected area) and rapid gradual shape recovery (Figure 1F-G). Such behavior was absent in the cells situated further away from the channel that were not reached by the calcium waves and served us as an internal control. Interestingly the temporal scale of nuclear shape recovery was closely correlated with the dynamics of calcium waves. We therefore probed if the observed process was dependent and regulated by the calcium bursts and wave propagation. Indeed, perturbation of calcium signaling with the known Ca²⁺ chelator BAPTA-AM^21^ perturbed the initiation and propagation of the calcium wave (Figure 1H-I) and abolished nuclear shape recovery (Figure 1J). We further confirmed the essential role of calcium on nuclear shape recovery by thapsigargin pretreatment that blocks the sarco/ER Ca²⁺ ATPase to empty ER calcium stores prior to folding^22^ (Figure 2C). Collectively these results showed that folding causes strong deformations of epithelial monolayer that are sufficient to generate calcium waves, which in response participate in the process of nuclear shape recovery.

**Figure 2.**
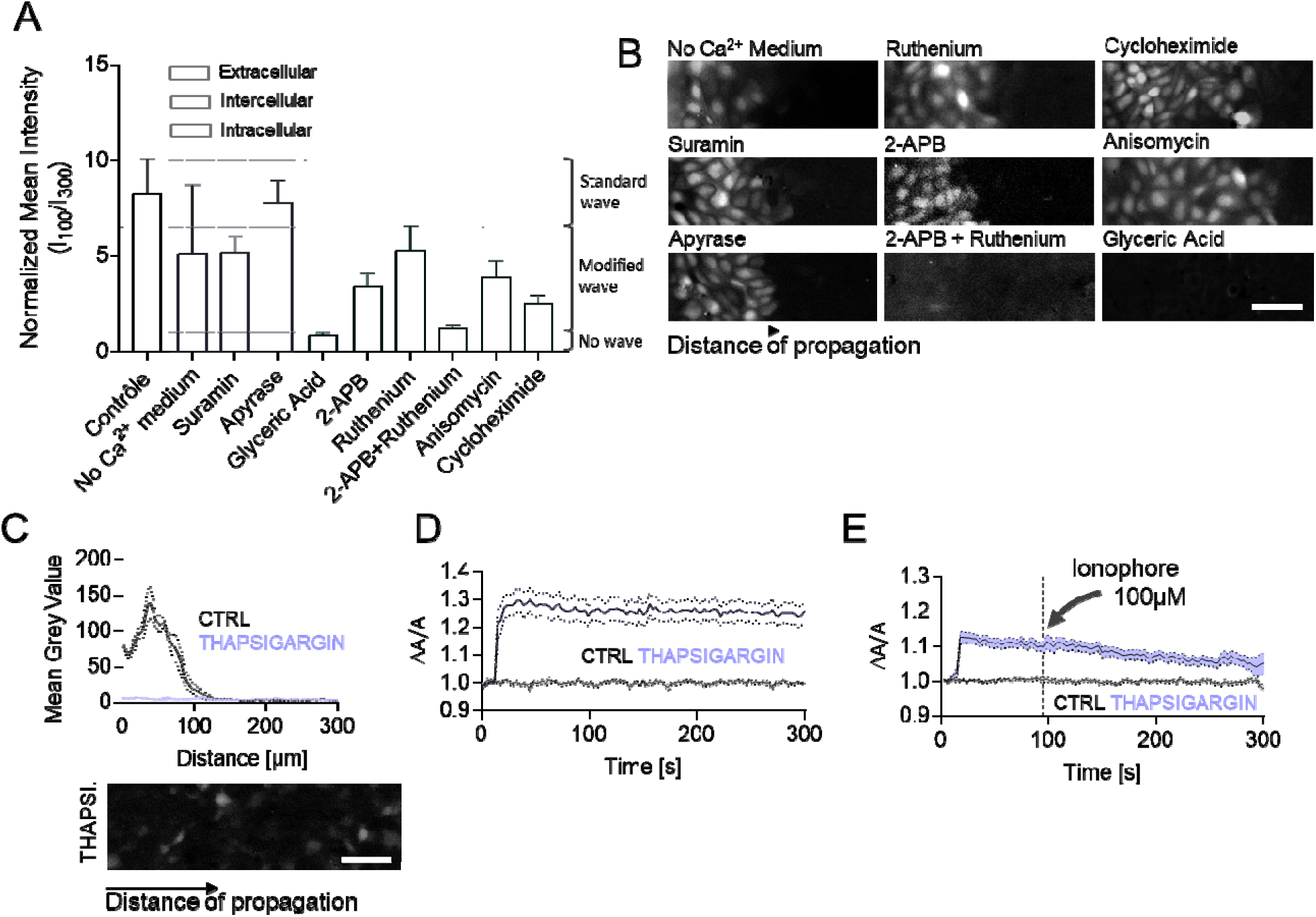
Intracellular and not extracellular calcium signaling is at the origin of calcium wave upon folding Calcium waves originate from the intracellular signaling and are necessary for nuclear recovery upon folding. A) Normalized mean intensity expressed as a ratio of intensity at distance 100um and 300 µm from the channel border as shown in the inset graph. Colors represent different conditions (including chemical inhibitors) targeting either extracellular, intracellular and intercellular calcium signaling. The ratio enable to determine the effect of treatment on the calcium wave intensity and propagation distance. Bar chart represents at least 10 measurements from 3 independent experiments; Error SEM. B) Images of Fluo4AM at t=T_max_ for control and treatments shown in A; Scale bar 50 µm C) Mean Grey value of Fluo4AM upon folding as a function of distance for control and Thapsigargin treated cells with the image representing the signal; Error bar: SEM, N=3 independent experiments; Scale bar 50 µm. D) Nuclear area changes in time upon folding for control and for Thapsigargin treated cells. Error bar: SEM, N=3 independent experiments pulled together E) Nuclear area changes in time upon folding for control and Thapsigargin treated cells. After 80 seconds following the fold the 100 µM Ionophore was added to the medium to induce extracellular calcium influx; Error bar: SEM, N=3 independent experiments.

### Intercellular and not extracellular calcium signaling is at the origin of calcium waves upon folding

To identify the pathway of calcium signaling we used a set of chemical inhibitors/activators to target pathways generally associated either with extracellular-, intracellular- or intercellular-signaling. Analysis of the effect of inhibitors was based on the creation of the Intensity as a function of distance at a given time point where calcium wave distance was maximal. The value of intensity at 100 µm from the channel border (I_100_) was divided by the intensity at 300 µm where typically calcium signal is unchanged (I_300_) (Figure 1D). The results are expressed as a ratio between I100/I300 and the classical calcium response resulted in the ratio superior to 7 (Figure 2A and 2B). We first investigated if calcium waves are associated with the calcium entry from the extracellular space. By using a medium without Ca²⁺ we showed that calcium waves are not induced by the extracellular calcium entry. In parallel, we investigated the role of extracellular ATP, which, in combination with its corresponding receptors on the cell surface (the ligand-gated P2X ion channels and a subset of G-protein-coupled P2Y receptors), mediates common autocrine and/or paracrine Ca²⁺ signaling mechanisms^23^. Cell pretreated with inhibitors of ATP with either Suramin^24^ or Apyrase^25^ did not show any significant difference as compared to control. ATP inhibition excluded mechanisms based on the extracellular paracrine signaling^26^.The second mechanism of calcium wave propagation is based on the transmission of ATP or Ca²⁺ molecules through the gap junctions which activate calcium rise in the following cells^27^. By using glyceric acid which blocks the gap junction transport^28^ we confirmed the intercellular nature of calcium wave propagation.

Next, we focused on intracellular pathways of calcium signaling, which target the endoplasmic reticulum. We pretreated cells with inhibitors perturbing major calcium receptors of the endoplasmic reticulum, the ryanodine receptor and the IP_3_ receptor. Treatment with 2-APB, a cell-permeable modulator of Ins(1,4,5)P3-induced Ca2+ release^29^ and with Ruthenium that targets ryanodine channels^30^ resulted in the calcium wave that propagating over shower distance and of the smaller intensity. The wave was fully abolished only under the combined treatment of 2-APB and Ruthenium. Interestingly, by pretreating cells with cycloheximide and anisomycin which both increase the residing concentration in the ER stores ^31,32^, resulted in calcium waves that have reached distance over 300 µm. Altogether our results show that the calcium source is intracellular (ER), but the wave propagation (from one cell to another) itself depends on intercellular signaling through gap junctions.

Finally, we wanted to test if the specific pathway of calcium signaling or the effective calcium concentration increase were necessary to trigger nuclear shape recovery. To do so, we used the fact that Thapsigargin pretreated cells did not trigger the calcium waves (Figure 2C) and thus lacked the response at the level of nuclei (Figure 2D). After 1.5 minutes post deformation to Thapsigargin pretreated cells, we added Ionophore^33^ (Figure 2E), which is known to internalize extracellular calcium. Interestingly, a simple rise in calcium was sufficient to activate nuclear shape recovery. Treatment of Ionophore alone on control monolayers did not trigger such response (Figure S1A). We were also able to activate nuclear shape recovery program by first pretreating cells with Thapsigargin followed by the activation of Piezo1 channel with Yoda activator^34^ (Figure S1B). Altogether, by combining pharmacological treatments we identified the source of Calcium concentration increase upon mechanical folding, and showed that calcium concentration is essential for the response at the nuclear level.

### Mechanism of Ca²⁺ propagation wave explained by a biokinetic model

To gain a deeper understanding of how the propagation of calcium waves is influenced by the cellular architecture, we have developed a reaction/diffusion model. As experiments showed no influence of extracellular Ca²⁺, our focus is solely on the dynamics of intercellular calcium. We assume that intercellular Ca²⁺ is transferred between cells via diffusion through gap junctions, as demonstrated by the previous experiments. The model relies on the one-dimensional, time-dependent reaction/diffusion equation, as described in Material and Methods. Our model predicts with good agreement to experimental results the concentration change in time for a cell at a given position with respect to the channel border (Figure 3A). Parameters’ values of the simulation, represented in the table Figure 3F are chosen in accordance with a good fit between experiments and numerical simulations. We validated the model by comparing the experimental data, where the duration of propagation from one cell to another is approximately 1.98 s (Fig1E), with the numerical data for which this duration is 1.9, expressed in dimensionless time (Fig3B). Thus, the conversion factor from dimensionless to real time is the ratio τ = 1.9 s. Knowing the mean experimental spacing between cells is L = 20 µm and the dimensionless diffusion coefficient found from the simulation is D* = 1, we can compute the real value of the diffusion coefficient in our model, as D = D*L²/ τ = 1 x 20²/1.9 = 2.1 x 10^-10^ m²/s, closer to the coefficient measured in *Xenopus laevis* oocytes of 2.2 x 10^-10^ m²/s ^35^.

**Figure 3.**
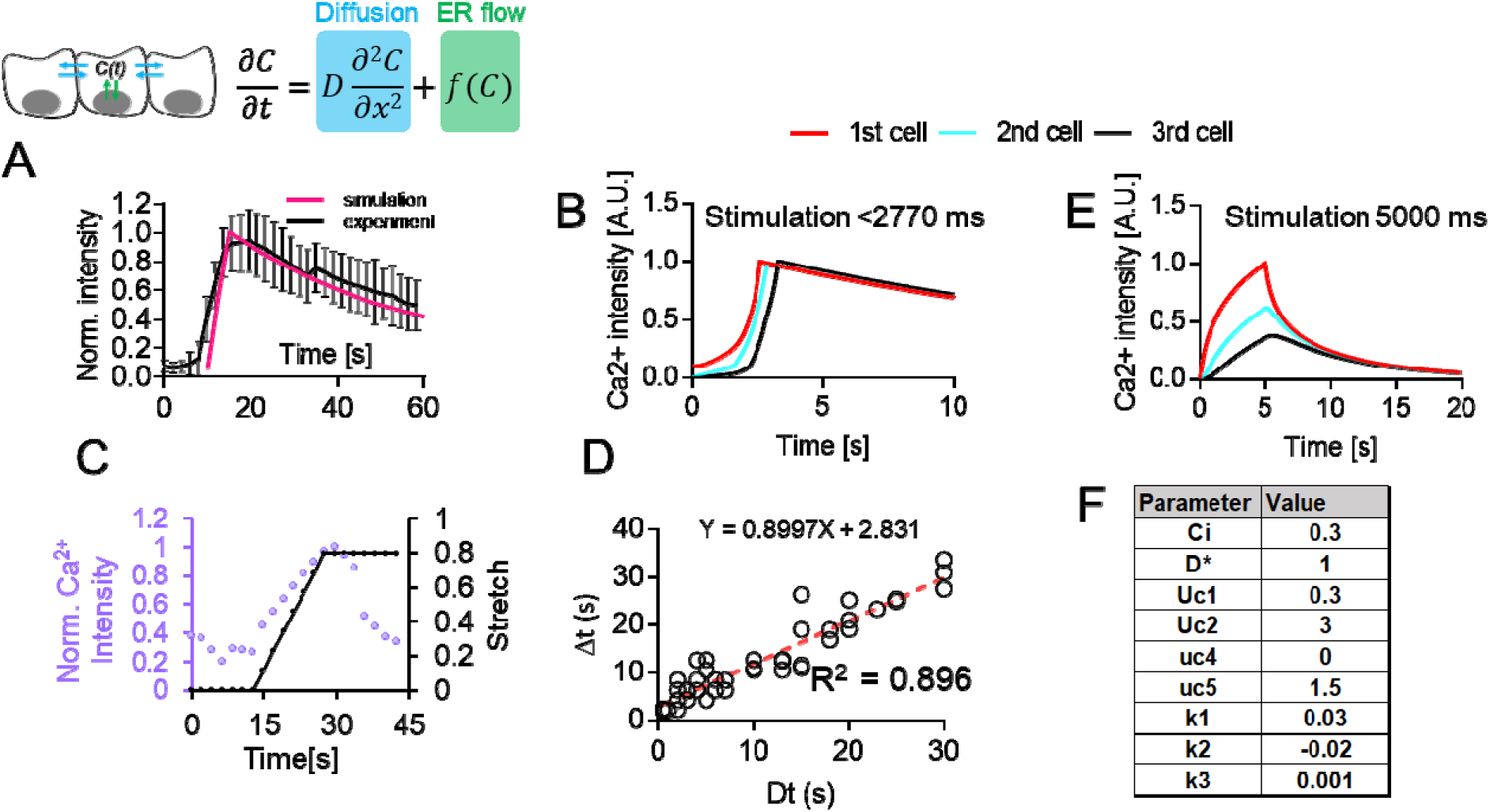
Mechanism of Ca²⁺ propagation wave explained by a biokinetic model. A) Numerical simulation to predict the wave profile in time based on ref and that takes into account the diffusion between the cells and the release of calcium from ER (as shown on the scheme). Red line represents the simulation and black the experiment. Error bar SD. B) Numerical simulation of calcium wave for 3 successive cells in a 1 dimensional lattice, in the case of a stimulation duration less than 2770 ms. Intensity peak of calcium are transferred from one cell to another over time. C) Changes in Fluo4AM signal (violet) as a function of time and the rate of stretch (black) for a single experiment. D) Proportionality between the rate of deformation with the rate of signal increase (represented in terms of increase durations Dt and Δt for numerous conditions. Each single point is an independent experiment E) Numerical simulation of calcium wave for 3 successive cells in a 1 dimensional lattice, in the case of a stimulation duration of 5000 ms. Intensity peak of calcium are not transferred from one cell to another over time. F) Table of values of the dimensionless parameters used for simulations presented in this figure.

Next, we wanted to interrogate experimentally and by modeling, how mechanical deformation rate regulates the calcium signaling. To do so, we generated folds by a progressive change in the pressure within the channel with simultaneous acquisition to follow intensity changes of Fluo4AM calcium concentration indicator (Figure 3C). Our results showed that calcium concentration increase was directly proportional (y=0.89x); the faster the cells were deformed the higher was the intensity of the calcium (Figure 3D). However, only rapid burst in calcium was triggering the calcium wave spreading. To test this further, we applied the progressive stretch in the model, by progressively increasing the concentration C in the cell 1, from 0 to C_i_, for a tuned duration Dt. As illustrated in Figure 3E, for duration of stimulation Dt = 5000 ms, wave propagation is not induced, since Ca²⁺ concentration is not conserved from cells to cells over time. However, for Dt < 2770 ms, the wave is propagated (Figure3B). This finding gives us an example on how tissues are sensitive to the duration of mechanical transduction as well as to the energy intensity involved, and explains why our previous experiments on progressive stretched may take too long to induce a Ca²⁺ wave. Altogether, these simulations confirmed experimental results described above and indicated implication of the ER as the source of the calcium and underlined the importance of strain rate on the generation of calcium waves.

### Level of nuclear envelope tension is essential to trigger nuclear shape recovery upon epithelial folding

To identify parameters that regulate nuclear shape recovery we tested if cellular density within the monolayer could change the rate or magnitude of the observed process. Interestingly, for highly dense monolayers we did not observe nuclear shape recovery despite the initial nuclear deformation (10^4^ cells/mm^2^ for highly confluent vs 2.8 10^3^ cells/mm^2^ for confluent) (Figure 4A). To understand how density of monolayers impacted the process we first verified if transport over gap junctions was perturbed. Western-blot analysis of Connexin-43 phosphorylation, which is present for permissive gap junctions ^36^, did not reveal any significant changes between the two investigated densities of cells (Figure 4B). Next, since nuclear lamina and nuclear stiffness determine the capacity of calcium release from ER^37^, we inspected changes in the nuclear lamina and nuclear shape by Lamin A immunofluorescence, to observe that highly confluent monolayers had nuclei significantly smaller and with many nuclear envelope folds (Figure 4C) suggesting that nuclear membrane tension was significantly smaller. To verify this, we used Flipper-TR technology combined with fluorescence lifetime imaging and compared tensional change at the plasma membrane and nuclear membrane for both cellular densities. Our results clearly indicated a significant drop of tension at the plasma and nuclear membrane (Figure 4D) for highly dense monolayers.

**Figure 4.**
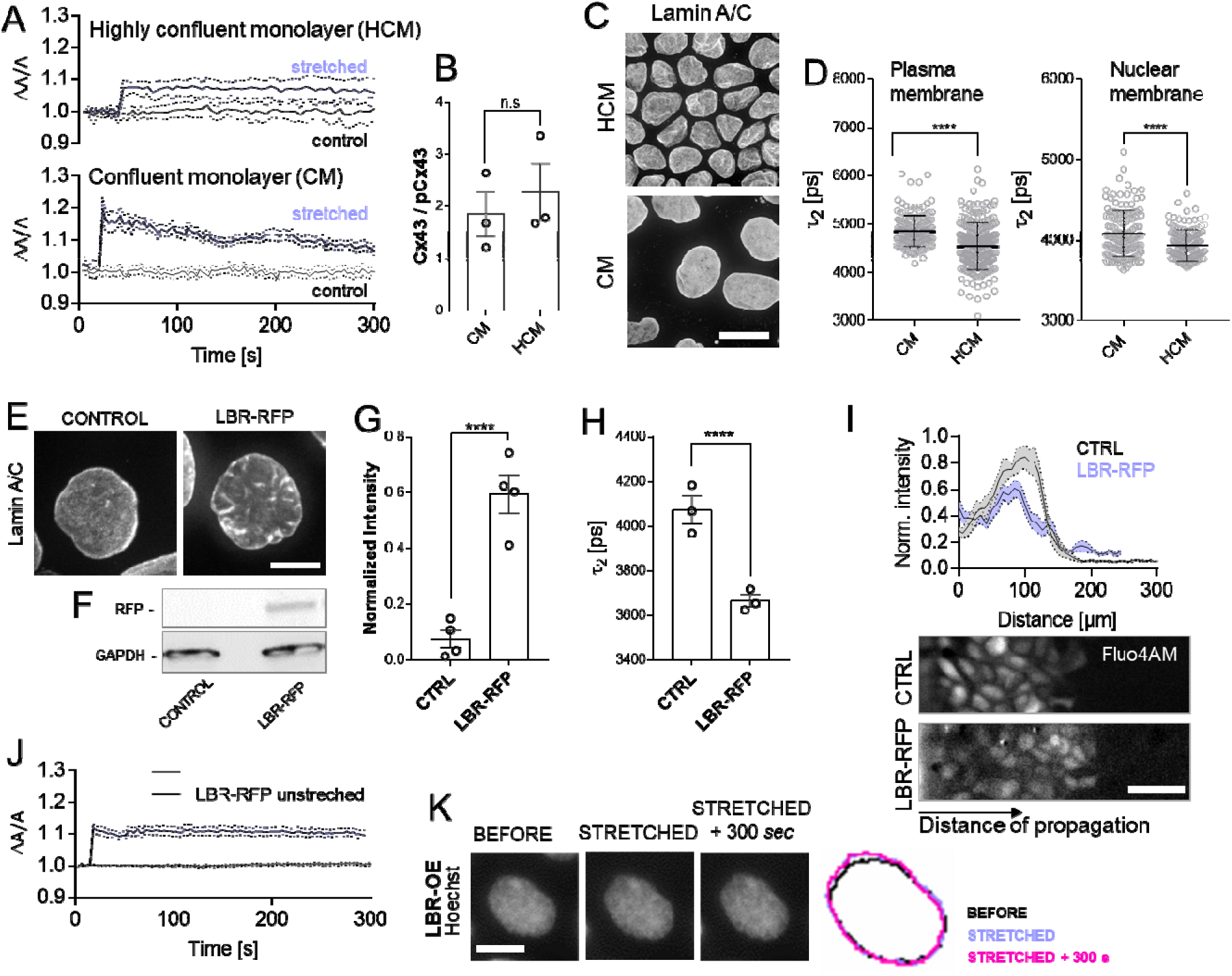
Level of nuclear envelope tension is essential to trigger nuclear recovery upon epithelial folding. A) Changes in nuclear area upon folding (violet) of confluent monolayers (CM) and highly confluent monolayers (HCM). Error bar: SEM, N=3 independent experiments B) Quantification of connexin43 phosphorylation state (Cx43/pCx43) based on the western blot experiments. Each point represents a separate replicate for CM and HCM; no statistically significant difference found, Mann-Whitney. C) Fluorescent images of CM and HCM stained with anti-Lamin A/C antibody; Scale bar: 10um D) Quantification of τ2 from Flipper-TR (plasma membrane) and ER Flipper-TR (nuclear membrane) stained cells using FLIM; n > 100 cells/condition pooled across three independent experiments; ****p <0.0001, Mann-Whitney; Error SEM. E) Fluorescent images with anti-Lamin A/C staining of control and cells with LBR-RFP overexpression; Scale bar 5 um F) Westernblot of control and cells overexpressing LBR-RFP with anti-RFP antibody. G) Quantification of LBR-RFP overexpression based on the westernblot from F. Each point represents an independent experiment. ****p<0.0001 Mann-Whitney. H) Quantification of τ2 from ER Flipper-TR (nuclear membrane) of control cells and LBR-RFP cells. N=3 Independent experiments; ****p <0.0001, Mann-Whitney; Error SEM. I) Normalized intensity profile of calcium wave (Fluo4AM) as a function of distance at t=T_max_ =18 s for control and LBR-RFP cells, and the corresponding images; Error SEM; Scale bar 50um J) Nuclear area changes of LBR-RFP cells upon folding in the function of time; Error SEM K) Images of single nucleus of LBR-RFP cells marked with Hoechst in three time points: before the stretch (t=6 s), stretched (t=15 s) and stretched + 300 s. The color masks of the nucleus correspond to different time points and come from the image analysis processing.

To understand the role of nuclear envelope tension we overexpressed LaminB receptor (LBR), which is known to decrease the tension at the nuclear membrane^38^. Overexpression of LBR of 40% as measured by western blot (Figure 4E-G) resulted in the appearance of wrinkled membrane even in low density cultures (Figure 3E). FLIM measurement of nuclear membrane tension using ER Flipper-TR tension probe confirmed that overexpression of LBR-OE resulted in the significant decrease of tension (Figure 3H). Interestingly, monolayers formed of these cells were capable to initiate calcium waves of similar magnitude and distance of propagation (Figure 4I). However, upon folding nuclei of LBR-OE failed to recover their shape (Figure 4J-K). Altogether, our results showed that two conditions need to be met to activate nuclear shape changes post folding: *i)* initiation and spreading of calcium waves (calcium concentration increase), and *ii)* deformation-associated nuclear envelope tension increase.

### Epithelial folding increases tension of nuclear envelope to activate cPLA2 and cell contractility

To identify the mechanism by which epithelial folding leads to nuclear shape recovery under the control of calcium wave and increase in tension, we first investigated the role of changes of chromatin state. In literature, changes in the in-plane tension resulted in chromatin de-condensation with the loss of H3K9me3 to mechano-protect cells from the DNA damage^39^. On the other hand, cellular flattening by axial confinement showed to increase chromatin condensation as observed by H3K9me3 signal increase^40^. To investigate if chromatin compaction is at the origin of nuclear shape changes, we performed H3K9me3 immunofluorescence on folded epithelial monolayers after 3 minutes post folding (Figure 5A). This specific time-scale matches the duration of the observed nuclear shape changes. Mean intensity analysis between the cells by the channel border and the internal control cells showed no significant difference between these conditions (Figure 5B). We observed also, that a part of population of nuclei expressed strikingly low levels of H3K9me3 as compared to other cells (indicated by arrows, Figure 5A), with higher percentage within the cells that were within the folded area (Figure 5C). Because in literature chromatin changes were inspected at 30 minutes post mechanical stress, we do not exclude the possibility of chromatin state changes at these longer times, but at the time-scale of the nuclear shape recovery this mechanism is unlikely to be involved, and therefore changes in nuclear membrane tension must activate another signaling pathway.

**Figure 5.**
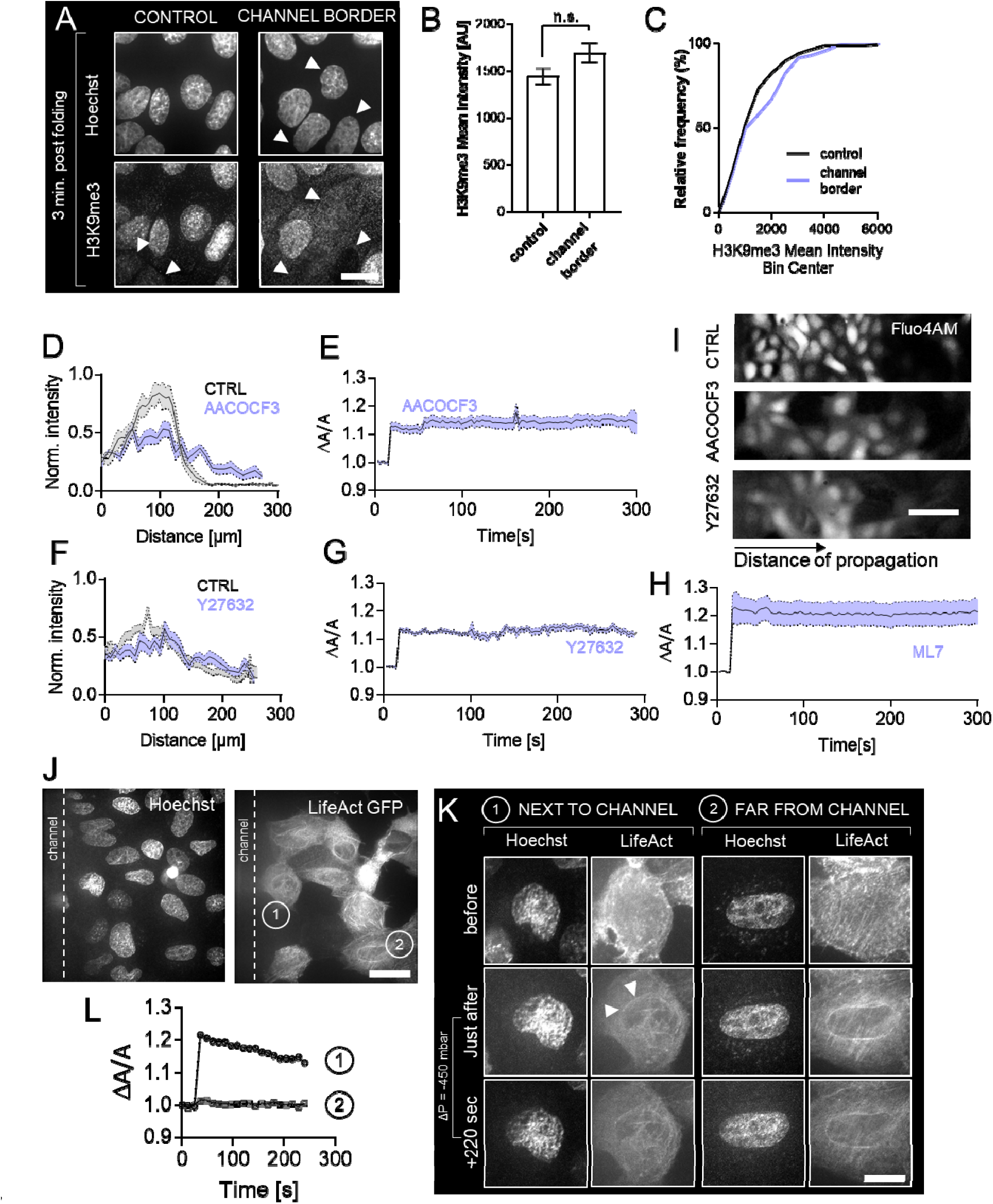
Epithelial folding increases nuclear tension which activates cPLA2 and cell contractility. A) H3K9me3 immunofluorescence images of control cells and cells by the channel border after 3 minutes post folding. Arrows indicate cells presenting low signal. Scale bar 10 µm. B) Quantification of the mean nuclear intensity of H3K9me3 staining. Error bar: SEM. N=3 independent experiments. X-test was performed for statistical significance. C) Relative frequency distribution represented in % of the H3K9me3 mean Intensity (From B) D) Normalized intensity profile of calcium wave (Fluo4AM) as a function of distance at t=T_max_=18s for control and cPLA2 inhibitor (AACOCF3) treated cells; Error SEM. E) Change in nuclear area upon folding (violet) as a function of time for AACOCF3 treatment. Error bar: SEM, N=3 independent experiments. F) Normalized intensity profile of calcium wave (Fluo4AM) as a function of distance at t=T_max_=18s for control and ROCK inhibitor (Y27632) treated cells; Error SEM. G) Change in nuclear area upon folding (violet) as a function of time for Y27632 treatment. Error bar: SEM, N=3 independent experiments. H) Change in nuclear area upon folding (violet) as a function of time for ML7 treatment. Error bar: SEM, N=3 independent experiments. I) Images of calcium wave for control, AACOCF3, and Y27632 treated cells; Scale bar 50um J) Images representing Hoechst stained nuclei and lifeact GFP actin organization at t=20 s post folding. 1 and 2 represent cells that were selected for the comparison in K and L; Scale bar 10 µm. K) Time-series of images of cell 1 and cell 2 showing the changes in the nuclear area and appearance of perinuclear actin rim upon folding. L) Representative time-evolution of nuclear area change post folding for example cells: 1 and 2 from J and K.

Activation of phospholipaseA2 pathway was observed under mechanical confinement^41^ or osmotic swelling ^42^, which led to nuclear membrane unfolding due to the rise of the tension. Previously, it was shown that activation of this nuclear phospholipase showed to activate the production of arachidonic acid (AA), which is a known regulator of myosin II activity ^43^. Therefore, to test this pathway we used an inhibitor of the cPLA2 activity, the AACOCF3^44^. We did not observe any significant effect of this inhibitor on the formation of calcium waves (Figure 5D and Figure 5I), yet the treatment perturbed the nuclear shape changes upon deformation (Figure 5E). Because cPLA2 pathway induces cellular contractility we tested if classical inhibitors of actomyosin contractility will also perturb the process of nuclear shape recovery. Pretreatment of monolayers with either ROCK kinase inhibitor (Y27632) (Figure 5F, 5G, and 5I) or with myosin light chain kinase inhibitor (ML7) (Figure 5H) confirmed that cell contractility was key to the nuclear shape recovery process.

We next inspected the response of actin cytoskeleton to tensional changes and calcium waves upon folding. In literature calcium bursts can promote actin flows with partial cytoskeleton depolymerization^45^ and formation of perinuclear actin rim^46^. Indeed, upon folding we observed partial actin depolymerization at the level of cortical actin, with basal actin fibers remaining intact (Figure S2A). Moreover, transient formation of the perinuclear actin rim was present in cells under tension as well in those that were reached by the propagating calcium wave (Figure 5J and 5K). Quantification of the nuclear projected area showed recovery only for cells that were under tension, and did not occur in cells positioned further away from the fold (Figure 5L) despite the presence of the nuclear actin rim. Additionally, measured high heterogeneity in the duration (Figure S2B) and the formation of the actin rim within the population of cells (Figure S2C) makes this transient structure unlikely to be directly involved in the observed nuclear retraction.

Collectively, our results identified two key elements that allow rapid intercellular communication and response orchestration upon mechanical deformation. Cells at the highest curvature within the fold are highly stretched, which triggers calcium waves and causes nuclear flattening (Figure 6B and Figure 6C). Changes of nuclear tension in the presence of calcium are essential to activate cPLA2 phospholipase signaling, which results in increased contractility in the population of cells within the fold, which is necessary to counteract the deformation induced by folding and at the scale of minutes rescues the nuclear deformation (Figure 6C).

**Figure 6.**
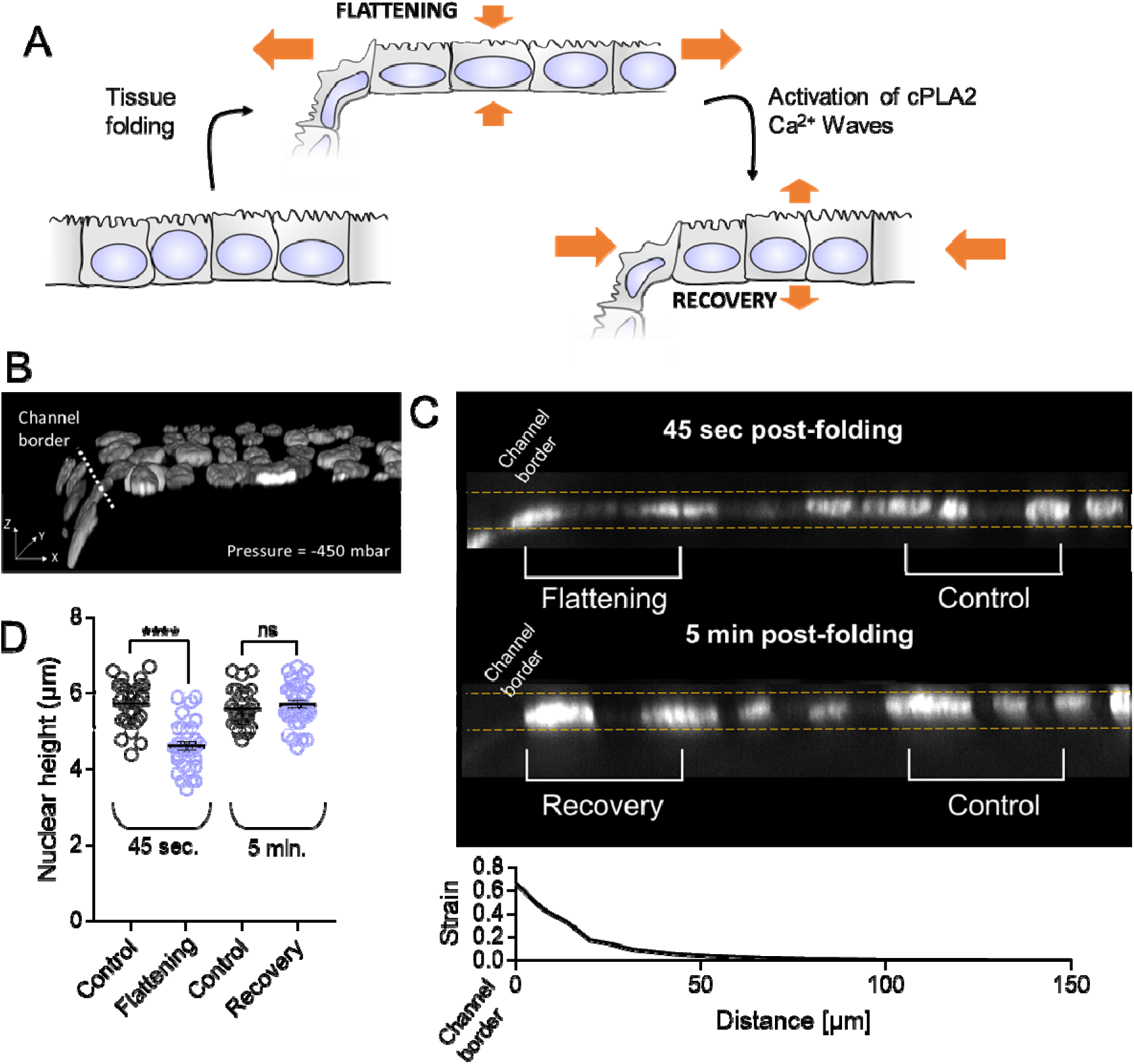
Proposed model of epithelial response and nuclear recovery upon folding. A) Scheme of the monolayer and nuclei shape changes upon stretching/folding and the subsequent response regulated by cPLA2 phospholipase and Ca2+ waves. Orange arrows in flattening indicate tissue stretching and in recovery cell contractility. B) 3D view of a confocal stack over the folded monolayer showing the change in the curvature at the channel border. Cells were fixed, stained with Hoechst and subsequently imaged. Confocal stack was obtained with the fold being maintained (ΔP = −450 mbar). Pointed line represents the border of the channel. Scale bar 10 µm. C) Single plane orthogonal view in the Z axis showing the resulting changes in the height of the nuclei that were stained with Hoechst. Channel border is situated on the left. Nuclei taken for analysis presented in D are defined (Flattening, Recovery and Controls). Graph below is in scale with the images and represents the spatial distribution of the strain starting from the channel border and outwards from the fold. D) Quantification of the nuclear height for monolayers folded and fixed/stained after 45 seconds and 5 minutes respectively. Nuclei were spatially selected as marked on C). Mann-Whitney test was used for statistical analysis. At least 26 nuclei were analysed per condition from N=2 independent systems for each time point. Error SEM.

## DISCUSSION

Epithelial tissues undergo extreme deformations during developmental processes and during their normal physiological function. This mechanical challenge requires cells to be equipped with specific programs that protect cells from mechanical damage and ensure tissue integrity. Lack of adaptation leads to the accumulation of mechanically induced DNA damage^39^ and can be at the origin of pathological states such as cancer. Mechanisms of tissue adaptation to mechanical stress function at different time-scales and only recently it has been shown that stretched epithelial monolayers can dissipate mechanical stress in the scale of the minutes ^47^, which is much faster than previously known mechanisms based on cell division, intercalation or extrusion^48^. In a model of stretched suspended epithelial monolayer, a biphasic stress relaxation has been observed with initial large-amplitude fast relaxation occurring during the first 6 seconds, followed by a smaller amplitude relaxation lasting about 2 minutes^47^. The time scale of the relaxation is comparable to observed rate of nuclear area recovery and slight increase of the characterisitic time can be associated with the fact that in our system monolayers are attached to a substrate. Indeed, another work that employed monolayer stress microscopy on pre-patterned epithelial islands, showed a rapid stress recovery in ∼80% within the first 2 minutes and remained stable until the stretch was released^49^. Moreover, in our previous study employing the same experimental system^20^, we showed that after 5 minutes from the fold induction, the relative tension level was increased, but its absolute value when estimated based on the calibration curve from^50^ was 10 fold lower than expected, confirming rapid stress relaxation. The mechanism of mechanical stress dissipation of stretched monolayers has been shown to be dependent on cellular contractility^47,49^. Whereas these works improve our understanding on the dynamics of stress dissipation in epithelia the mechano-transduction pathways that trigger specific active response to stress and the role of the nucleus in this response is missing.

In our study, we complement the current state of knowledge by providing mechanistic description on how local mechanical stress and curvature regulate nuclear shape and orchestrate relaxation. Assuming that nuclear volume is maintained^51,52^ we showed that epithelial folding resulted in significant nuclear flattening that is recovered linearly at the time scale of minutes and is blocked by contractility chemical inhibitors. Intrestingly, whereas nuclear area recovered partially in response to folding, the nuclear height returned to its control level. This suggested that besides of contractility based recovery, an osmotic swelling can be potentially involed as previously showed in folded epithelia^53^. The first essential component for relaxation that we identified is the calcium wave that spread across the epithelium outward from the triggered folds. Through mathematical modelling and experimental use of chemical inhibitors of calcium receptors/channels we identified that the calcium burst comes from the ER, and the calcium waves propagate through gap junctions. Interestingly, similar behavior of nuclear retraction regulated by Piezo1 activity and associated to calcium signaling has been previously observed in cells exposed to shear stress^54^ pointing out a different mechanism of calcium release. From that perspective, experiments with Thapsigargin and Ionophore employing extracellular calcium proved that nuclear retraction occurs under a strong calcium burst but the source of this burst is indifferent for the process which underlies the universality of the calcium mechano-transduction signaling at play. Despite the capacity of calcium to condense chromatin^55^ and potentially impact nuclear shape, we showed that at such short time scale chromatin remodeling is minimal. We further showed that calcium burst alone is insufficient to trigger nuclear shape changes and a second condition must be met. Our experiments on epithelia composed of cells having normal plasma membrane tension but relaxed nuclear envelope (LBR overexpression as in ^41^), confirmed the importance of nuclear tension change (occurring due to passive flattening) to trigger stress relaxation. We identified that cPLA2 pathway activation associated with the nuclear tension increase is at the origin of the observed process and coordinates the response at the tissue scale level. This pathway has been previously involved in wound healing processes^42^ and in activating single cell amoeboid migration^41^ showing that nuclear tension changes regulate cell function and tissue homeostasis.

Our previous work showed that depending on the direction of the folding the functional response of the cells was different. On folds towards basal membrane with increase of tension we observed calcium waves but no transcriptional changes, whereas opposite folding associated with the negative tension prevented calcium waves formation but promoted a strong signature at the gene expression level^20^. This is surprising, as numerous works showed that positive tension (stretch) regulates transcription^56,57^. Yet, as we show here, in the local epithelial deformation these nuclear shape changes are rapidly recovered. From that perspective, it is possible that described here nuclear recovery can serve as a fast mechano-protective mechanism ensuring transcriptional stability upon dynamically occurring tissue shape changes in development and in during physiological processes.

Altogether, our work identified nucleus as a coordinator of collective cell response to local epithelial folding. We showed that folding and associated changes in curvature and tension trigger calcium waves that together with changes in nuclear envelope tension activate cPLA2 pathway. In our system where nuclear deformation occurred instantly upon folding, cPLA2 is essential for relaxation associated nuclear shape recovery that occurred in the scale of minutes post folding. Therefore, **based on the identified mechanism we propose that nucleus is the internal sensor of mechanical stress and triggers the signaling cascade leading to actomyosin contractility serving to dissipate the stress.**

## Authors contributions

M.B.-C.-B. and M.E.D. performed all experiments. L.P. and Ch.O. assisted in the establishment of stable cell lines and molecular biology experiments. K.E. participated in the nuclear retraction quantification experiments both in vitro and in vivo. C.G. partially financed M.B.-C.-B and the project. All authors contributed to the discussion and creation of the manuscript.

## Acknowledgements

We would like to thank the MicroCell microscopy facility for their time, discussions, and expertise. We would especially like to thank Mylene Pezet for the FACS sorting of our cells for stable cell line establishment. We also thank Isabelle Tardieux and her team for allowing us to perform imaging in her laboratory. The project and M.B.-C.-B. were financed by the ANR funding agency (CellCOMM ANR-17-CE13-0006).

---

**Figure S1.**
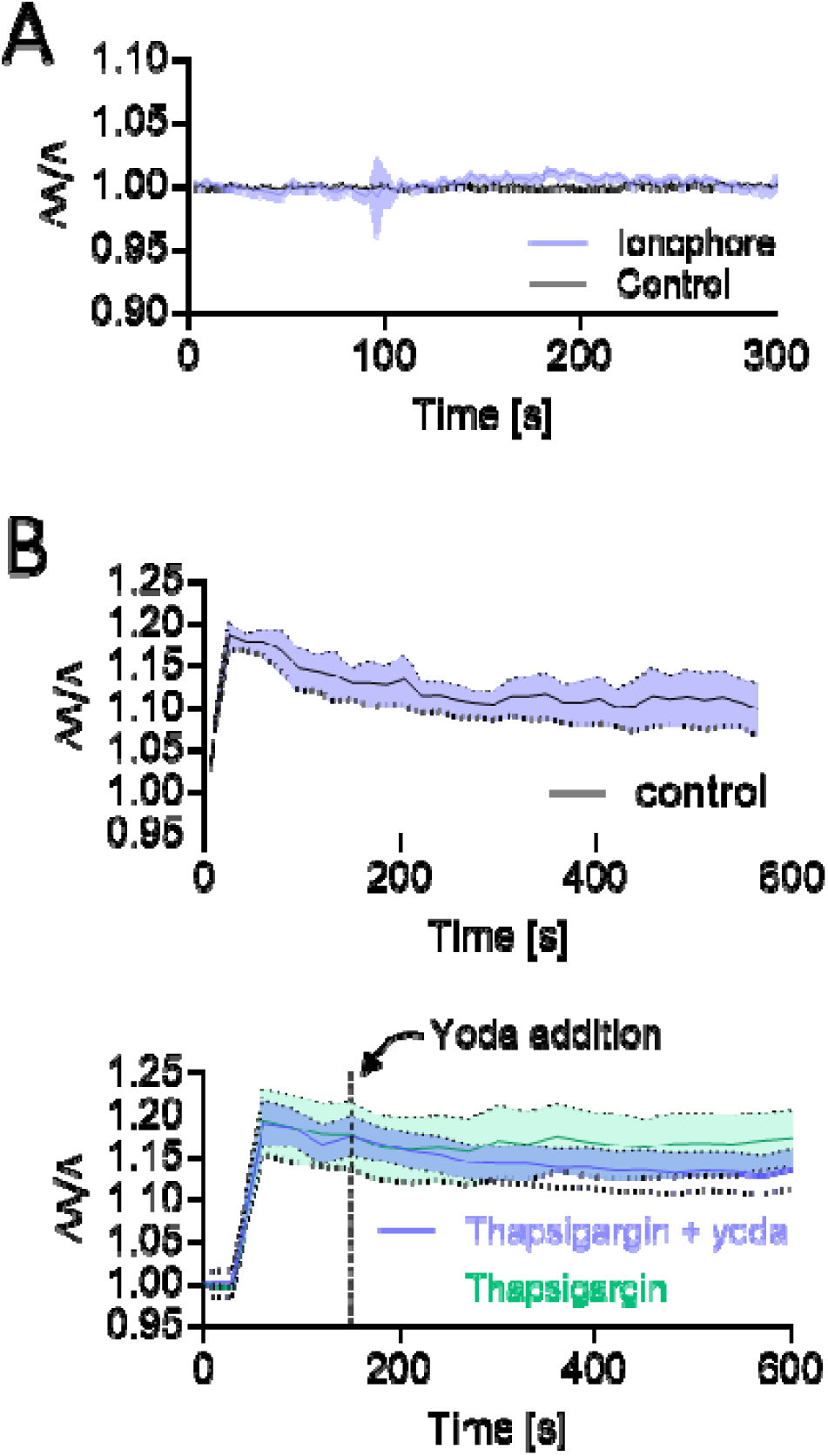
A) Effect of addition of Ionophore on calcium levels as compared to non-treated cells (control). Error bar: SEM, N=3 independent experiments. B) Evolution of nuclear projected area in time in control cells, and in cells treated with either Thapsigargin or Thapsigargin with addition of Yoda after 150 s following the induced folds. Error bar: SEM, N=3 independent experiments.

**Figure S2.**
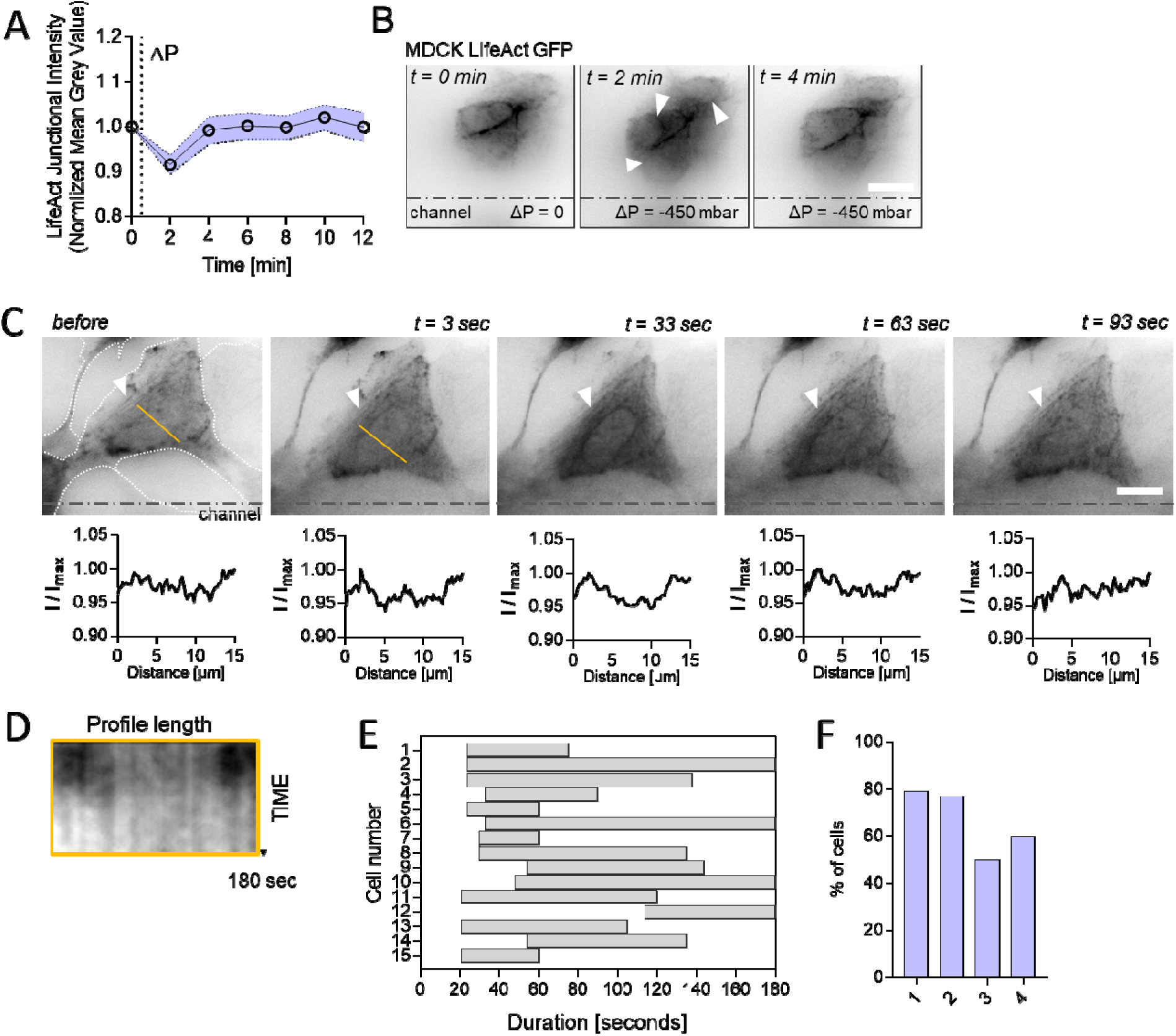
A) Change of actin intensity in time as measured over the cell-cell junctions. Pointed line indicates the moment of fold induction. Error bar: SEM B) Images of MDCK LifeACT GFP cells showing the occurring changes in the intensity of junctional/cortical actin over time before (t=0) and post folding. Scale bar 10 µm. C) Time-serie of a LifeAct GFP cell position by the channel border and within the monolayer (borders of other cells are marked with the pointed line). Orange line indicates the profile over which intensity was ploted (below the images). Example of actin fibers that remained intact is marked by the white triangle. Scale bar 10um. D) Kymograph over the profile marked in (C) showing the transient appearance of nuclear actin rim. E) Measured duration of existence of the actin rim among the population of cells. N=3 independent experiments. F) Measurement of % of cells by the channel border that formed perinuclear actin rim. 1-4 represent independent experiments.

**Table S1.** Chemical reagents used in the study.

| Reagent | Function | C (mM) | Duration | References |
| --- | --- | --- | --- | --- |
| Ruthenium | Ryanodin receptor inhib. | 10 | 40 mn | Merck #557450-250MG "Ruthenium Red - CAS 11103-72-3 - Calbiochem" |
| Glycyrrhetic Acid | Gap junction inhib. | 10 | 30 mn | Merck #G8503 "18α-Glycyrrhetic acid" |
| Thapsigargin | Serca pump inhib. | 10 | 45 mn | Merck #SML1845 "Thapsigargin Ready Made Solution" |
| BAPTAM | Ca<sup>2+</sup> chelator in cytosol |  | 1 h 30 mn | Merck #A1076-25MG "BAPTA-AM" |
| AACOCF3 | CPLA2A phospholipase inhib. | 14 | 15 mn | Biotechne #1462 "AACOCF3" |
| Ionophore Ca<sup>2+</sup> | Permeate plsm. membrane to Ca<sup>2+</sup> | 10 | instant | Merck #C7522 "Calcium Ionophore A23187" |
| IP3 | Activate IP3 receptor in ER membrane | 1 | 30 mn | Merck #I9766 "IP3 (D-myo-Inositol 1,4,5-tris-phosphate trisodium salt)" |
| Suramin | Purenergic receptor inhib. | 10 | 40 mn | Santa Cruz #sc-507209 "Suramin sodium - 50 mg" |
| Yoda | Open Piezo 1 channel | 25 | 60 mn | Merck #SML1558 "Yoda1" |
| Anisomycin | Block translocon translation on ER | 6.55 | 30 mn | Merck #A9789-5MG "Anisomycin from Streptomyces griseolus" |
| 2-APB | IP3 receptor inhib. | 10 | 40 mn | Abcam #ab120124 "2-APB, Ca<sup>2+</sup> release modulator" |
| Cycloheximide | Block translocon translation on ER | 10 | 2 h | Merck #C4859-1ML "Cycloheximide solution" |
| Fluo4AM Direct | Ca<sup>2+</sup> live probe in cytosol, Green | x2 | 40 mn | Invitrogen #F10471 "Fluo-4 Direct™ Calcium Assay Kit" |
| Flipper-TR | ER probe, Blue White | 1μL/mL | 1 h 30 mn | Invitrogen #E12353 "ER-Tracker™ Blue-White DPX, for live-cell imaging" |
| ER FlipperTR | ER probe, Blue White | 1μL/mL | 1 h 30 mn | Invitrogen #E12353 "ER-Tracker™ Blue-White DPX, for live-cell imaging" |
| ML7 | Myosin light chain kinase inhib | 10 | 1 h | Merck #I2764 "ML7" |

## Materials and Methods

### Cell culture

MDCK II cells (Merck, #00062107-1VL) and MDCK LifeAct GFP (gift from Dr Destaing), and LBR-RFP were cultured in MEM (Life Technologies) supplemented with 5% fetal bovine serum (Sigma) and 1% penicillin/streptomycin (Dutscher). Subculturing occurred every 3 days at approximately 70% confluence using trypsin-EDTA (Dutscher). To establish a dense, quiescent monolayer of MDCK cells, we employed a calcium switch method following a previously described protocol (Benham-Pyle et al., 2016).

For the generation of stable cell lines overexpressing LBR, viruses were generated in HEK 293T cells in a 100-mm dish by simultaneous transfection of 5 µg psPAX2 packaging vector, 2.5 µg pVSVG envelope plasmid, and 7.5 µg pSico-tagRFP-LBR vector (Ozyme #632587). Transfection media were replaced after 4 hours with 5 ml of complete medium. Supernatant containing lentiviral particles was collected 48 hours post-transfection and filtered through a 0.45-µm syringe filter. MDCK cells seeded in a six-well plate were transduced with 0.5 to 1 ml of viral particles in 2.5 ml of complete. Following 48 hours of culture, cells were transferred to T25 flasks and cultured for 2 weeks. Then, RFP-tagged cells were subsequently isolated via FACS and seeded for amplification.

### Drug Treatment

Drug treatments were conducted with at least three independent experiments. Prior to stretch and calcium wave video acquisition, cells were incubated with reagents at a specific concentration and duration (see *Table 1* in supplementary information)

### Plasmid construction

Plasmid encoding LBR was constructed using a psicoR backbone for subsequent lentiviral transfection. Standard plasmid and LBR sequences were obtained from Addgene (ref:61996, LBRpEGFP-N2). Subsequently, LBR sequence was cloned and fused with a tagRFP sequence. This sequence fusion was then inserted into a PSICO plasmid downstream of a CAG promoter.

### Flow cytometry

The cells were subjected to flow cytometry analysis using a jet-in-air cell sorter (AriaIIU, BD-Biosciences). tagRFP fluorescence was excited by a 561-nm laser and collected through a 600 LP filter and 610/20 emission filters. The cell suspension was passed through a 100-micron nozzle at a sheath pressure of 20 psi and a drop-drive frequency of 30 MHz, with a flow rate set at 2 and a threshold rate around 500 events/s. The purity sort mode was selected for sorting, and tagRFP positive cells were collected in a 1.5-ml Eppendorf tube containing 100 µl of MEM.

Sorting gates were established using a negative control (non-tagRFP cells) and a positive control (cells uniformly expressing tagRFP. The sorted fraction (tagRFP) was subsequently cultured for amplification.

### Microfabrication

The microfluidic devices were produced by directly milling the channel structure into transparent, 375-µm-thick polycarbonate film (Lexan 8010, Goodfellow Cambridge) using a vertical CNC milling machine (MFG4025P, Ergwind, Poland) and a 2-flute fish-tail endmill cutter with diameters of 400 or 200 µm (FR208, InGraph, Poland). The milled channels were cleaned with a pressure washer (K7 Premium, Karcher, Germany) to eliminate any remaining swarf and loosely bound bulk material formed during milling. Subsequently, the milled chips were manually washed with isopropanol and deionized water and then dried with compressed air. Polycarbonate slabs were bonded with a PDMS membrane of 20 µm thickness (Wacker, ELASTOSIL Film 2030 250/20) using 1% 3-aminopropyltriethoxysilane (APTES, Merck) following a previously described method (Sunkara et al., 2011).

In brief, clean polycarbonate slabs underwent 1-minute exposure to oxygen plasma (Diener, 100% power, MHz generator) and were subsequently incubated with an APTES solution for 20 minutes. After thorough washing with dH20 and isopropanol, the polycarbonate was firmly pressed into contact with the PDMS membrane freshly activated by oxygen plasma (1 minute, 100% power). Bonding occurred within minutes thereafter.

### Immunofluorescence

The cells were initially fixed with methanol (Sigma) for 6 minutes at −20°C, then permeabilized using 0.1% Triton in PBS (Sigma), followed by washing with PBS and blocking with a solution consisting of 2.5% bovine serum albumin in PBS Tween 0.2% for 1 hour. Subsequently, the samples were incubated overnight at 4°C with primary antibodies (H3K9me3 abcam #ab8898, connexin43 thermofisher #13-8300, and p-connexin43 thermofisher #PA5-104820) in the blocking solution, followed by three washes with PBS Tween 0.2%. The cells were then incubated with secondary antibodies at room temperature for 1 hour, followed by three additional washes with PBS Tween 0.2%. After two washes with PBS, the samples were mounted using mounting medium with DAPI (Prolong—Invitrogen). For nuclear height analysis, monolayer with triggered fold were fixed on the microscope, stained with Hoechst and mounted with the mounting medium (as described above). Stained systems then were inverted and imaging occurred through the mounted coverslip with the pressure mainted in the channel throughout the procedure.

Cells were imaged using a Spinning Disk Andromeda (iMIC 2.0, TILL photonics-FEI) with an EMCCD iXon 897 Camera and an alpha-Plan Apo 63×/1.46 oil objective.

### Western Blotting

Standard Western blotting protocols were utilized. Cells were lysed at 4°C directly in Laemmli buffer for Western blotting, which consisted of Tris–HCl pH 6.8 (0.12 M), glycerol (10%), sodium dodecyl sulfate (5%), β-mercaptoethanol (2.5%), and bromophenol blue (0.005%).

Lysates were applied onto NuPAGE 4–12% 15-well Bis-Tris gels (Invitrogen; NP0323BOX), NuPAGE™ 10% Bis-Tris, 1.0 mm, 10-well protein gels (ThermoFisher Scientific; NP0301BOX), and subjected to electrophoresis at 150 V for 1 hour. The gels were then wet-transferred using 70% ethanol onto Nitrocellulose membranes sized μM; 7 cm × 8.5 cm (P/N 926-31090). Subsequently, the membranes were blocked for one hour at room temperature using BSA. Primary antibodies were diluted in the blocking buffer and incubated overnight at 4 °C. Following this, washes were carried out with TBS 0.1% Tween-20 (TBST) before applying the secondary antibody for one hour at room temperature. Further washes were performed with 1× TBS prior to imaging on Vilber imaging system.

### Video microscopy

Live cells were observed using the AxioObserver inverted stand (Carl Zeiss) equipped with 40× and 20× objectives [LD Plan-Neofluar 20X N.A. 0.4 Ph2 Korr bague 0-1.5, and UPlanFl LD water 40X N.A. 1.1) and an on-stage cell incubation chamber. Epifluorescence were achieved using the HXP-120 (Carl Zeiss) metal-halide fluorescence light source connected via a liquid light guide. ND attenuation filters were utilized to adjust the excitation intensity to 50%, 25%, and 10% of maximum. Fluorescence imaging was performed with a cooled MRm3 (Carl Zeiss) camera at 0.5 image/s. Live nuclear deformation measurements were performed using ImageJ.

### Image analysis

Time-lapse images of calcium wave propagation were processed using Fiji software. For each video, the background was removed by computing the mean of every pixel in the video before stretching creating a mean background image, then by subtracting each image of the video by the mean image. Then, several bands of cells were segmented. To analyze the wave spatially, co-linear bands were segmented, and Fluo-4 AM fluorescence intensity, indicative of Ca²⁺ levels, was quantified using the “Plot profile” function (as in ^58^).

For temporal analysis, bands orthogonal to the direction of propagation were segmented, and intensity was quantified using the “Z-stack” function. Multiple band positions relative to the mechanically stressed area were segmented to accurately describe propagation dynamics. For each experiment, data were normalized by dividing by the maximum intensity value of Fluo4am over time or along the x-axis. Images of nucleus retraction were analyzed using Fiji software. Each nucleus, labeled with Hoechst, was segmented, and their areas were tracked using the “Analyze Particle” function for quantification. For each experiment and each nucleus, data of its area were normalized by dividing its mean area values nucleus before stretching.

For nuclear height quantification, nuclei were cut out from the total image at the region by the channel to measure flattening/recovery and away from channel as the internal controls. Single nuclei stacks were resliced to present the Z view followed by maximal projection. On created Z-view, threshold was used to define the nuclei shape in Z-axis, and rectangle function was fitted to measure the maximal height of the nuclei.

### FLIM Microscopy

Flipper-TR fluorescent tension probe (Spirochrome) was used according to the provided protocol in order to quantify plasma membrane tension. Briefly, we incubated a confluent epithelial layer with 1 mM Flipper-TR (ER Flipper-Tr (#SC021) or Flipper-TR (#SC020); 1µM) probe during 1-2 hours. FLIM imaging was performed using a Zeiss LSM 710 confocal microscope equipped with a time correlated single photon counting module from Becker&Hickl GmbH. SPCM software was used to record the data in photon-counting mode using a 40x/1.1 C-Apochromat water objective (600 mm working distance). Excitation was performed using a pulsed two photon laser (Ultra-II, Coherent) at 900 nm with emission collected through band pass 575/610 nm filter gated with a hybrid GaAsP-APD detector. For each image signal was accu-mulated during 180 seconds. SPCImage software was used to fit fluorescence decay data to a dual exponential model. Incomplete multi-exponentials model was used to take into account fast repetition rate of the laser (80 MHz). We quantified decay time τ_2_ (of regions corresponding to intercellular junctions or nuclear membrane) using ImageJ by averaging τ along the manually selected ROI over the cell-cell interface or at the nuclear membrane.

### Numerical simulations

We are investigating the behavior of endothelial cell networks, which are composed of one-dimensional chains of cells and the interconnected networks they form. To better understand how calcium wave propagation is affected by cellular architecture, we have developed a reaction-diffusion model. Experimental data indicate that extracellular Ca²⁺ has a negligible impact, so our study focuses exclusively on intercellular calcium dynamics. We assume that intercellular Ca²⁺ transfer occurs through diffusion via gap junctions, as suggested by prior findings. The model is based on a one-dimensional, time-dependent reaction-diffusion model, taken from^59^:

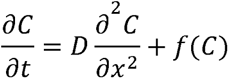

Here, *C* represents cytosolic calcium concentration, *D* denotes diffusion between cells via gap junctions, and *f* represents the rate of intracellular calcium concentration change. Calcium dynamics are influenced by the ER, which releases or absorbs calcium through IP_3_ and Ryanodine receptors. This leads to calcium-induced calcium release (CICR), a self-amplifying process controlled by thresholds UC1 and UC2 modeled by:

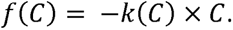

Below the UC1 value, calcium is absorbed by the ER so *k*(*C*) = *k*1 > 0; between UC1 and UC2, it is released and *k*(*C*) = *k*2 < 0; and above UC2, calcium is either reabsorbed or expelled i.e. *k*(*C*) = *k*3 > 0. The model incorporates finite-difference discretization to simulate calcium diffusion across a multicellular chain, assuming constant diffusion coefficients and no signal attenuation. It captures key features of intercellular calcium signaling observed experimentally.

Exclusively in our study, the diffusion coefficient D is no longer constant. A threshold value has been introduced concerning the concentration *C*, such as *D* = *D*(*C*). Namely, *D*(*C*) = 0 if C < *C*\* and *D*(*C*) = *D*\* if *C* > *C*\*. We decided to keep *C*_i_, the initial concentration introduced in the cell 1, for the next experiment.

For implementing a progressive stretch, a progressive increase of Ca²⁺ was added in the cell 1 until from *C* = 0 to *C* = *C_i_*, during *Dt*.

### Statistics

Statistical analysis was conducted using Prism Software. Data are expressed as stated by mean ± standard deviation error of the mean (SEM) or standard deviation (SD). Unless otherwise specified, unpaired t-tests with respective corrections were employed. The number of independent experiments conducted for all quantitative data is specified in the figure legends.

## References

1. Rauzi, M. et al. Embryo-scale tissue mechanics during Drosophila gastrulation movements. Nat Commun 6, 8677 (2015).

2. Davidson, L. A. Epithelial machines that shape the embryo. Trends Cell Biol 22, 82–87 (2012).

3. Polyakov, O. et al. Passive mechanical forces control cell-shape change during Drosophila ventral furrow formation. Biophys J 107, 998–1010 (2014).

4. Falo-Sanjuan, J. & Bray, S. J. Notch-dependent and -independent transcription are modulated by tissue movements at gastrulation. Elife 11, (2022).

5. Rauzi, M., Hočevar Brezavšček, A., Ziherl, P. & Leptin, M. Physical models of mesoderm invagination in Drosophila embryo. Biophys J 105, 3–10 (2013).

6. Dolmetsch, R. E., Xu, K. & Lewis, R. S. Calcium oscillations increase the efficiency and specificity of gene expression. Nature 392, 933–936 (1998).

7. Machaca, K. Ca2+ signaling, genes and the cell cycle. Cell Calcium 48, 243 (2010).

8. Bhosale, G., Sharpe, J. A., Sundier, S. Y. & Duchen, M. R. Calcium signaling as a mediator of cell energy demand and a trigger to cell death. Ann N Y Acad Sci 1350, 107–116 (2015).

9. Haws, H. J., McNeil, M. A. & Hansen, M. D. H. Control of cell mechanics by RhoA and calcium fluxes during epithelial scattering. Tissue Barriers 4, 1–15 (2016).

10. Dupont, G., Combettes, L., Bird, G. S. & Putney, J. W. Calcium oscillations. Cold Spring Harb Perspect Biol 3, 1–18 (2011).

11. Leybaert, L. & Sanderson, M. J. Intercellular Ca2+ waves: Mechanisms and function. Physiol Rev 92, 1359–1392 (2012).

12. Smedler, E. & Uhlén, P. Frequency decoding of calcium oscillations. Biochim Biophys Acta 1840, 964–969 (2014).

13. Peyronnet, R., Tran, D., Girault, T. & Frachisse, J. M. Mechanosensitive channels: Feeling tension in a world under pressure. Front Plant Sci 5, 1–14 (2014).

14. Lane, M. C., Koehl, M. A. R., Wilt, F. & Keller, R. A role for regulated secretion of apical extracellular matrix during epithelial invagination in the sea urchin. Development 117, 1049–1060 (1993).

15. Miller, A. L., Fluck, R. A., McLaughlin, J. A. & Jaffe, L. F. Calcium buffer injections inhibit cytokinesis in Xenopus eggs. J Cell Sci 106 (Pt 2), 523–534 (1993).

16. Chang, D. C. & Meng, C. A localized elevation of cytosolic free calcium is associated with cytokinesis in the zebrafish embryo. J Cell Biol 131, 1539–1545 (1995).

17. Webb, S. E., Lee, K. W., Karplus, E. & Miller, A. L. Localized calcium transients accompany furrow positioning, propagation, and deepening during the early cleavage period of zebrafish embryos. Dev Biol 192, 78–92 (1997).

18. Antunes, M., Pereira, T., Cordeiro, J. V., Almeida, L. & Jacinto, A. Coordinated waves of actomyosin flow and apical cell constriction immediately after wounding. J Cell Biol 202, 365 (2013).

19. Brun-Cosme-Bruny, M. et al. Microfluidic system for in vitro epithelial folding and calcium waves induction. STAR Protoc 3, 101683 (2022).

20. Blonski, S. et al. Direction of epithelial folding defines impact of mechanical forces on epithelial state. Dev Cell 56, 3222–3234.e6 (2021).

21. Collatz, M. B., Rüdel, R. & Brinkmeier, H. Intracellular calcium chelator BAPTA protects cells against toxic calcium overload but also alters physiological calcium responses. Cell Calcium 21, 453–459 (1997).

22. Gerasimenko, J., Maruyama, Y., Tepikin, A., Petersen, O. H. & Gerasimenko, O. Calcium signalling in and around the nuclear envelope. Biochem Soc Trans 31, 76–78 (2003).

23. Schwiebert, E. M. Extracellular ATP-mediated propagation of Ca2+ waves. Focus on ‘mechanical strain-induced Ca2+ waves are propagated via ATP release and purinergic receptor activation’. Am J Physiol Cell Physiol 279, (2000).

24. Calcaterra, N. B., Vicario, L. R. & Roveri, O. A. Inhibition by suramin of mitochondrial ATP synthesis. Biochem Pharmacol 37, 2521–2527 (1988).

25. Komoszyński, M. & Wojtczak, A. Apyrases (ATP diphosphohydrolases, EC 3.6.1.5): function and relationship to ATPases. Biochim Biophys Acta 1310, 233–241 (1996).

26. Kawano, S. et al. ATP autocrine/paracrine signaling induces calcium oscillations and NFAT activation in human mesenchymal stem cells. Cell Calcium 39, 313–324 (2006).

27. Benninger, R. K. P., Zhang, M., Steven Head, W., Satin, L. S. & Piston, D. W. Gap junction coupling and calcium waves in the pancreatic islet. Biophys J 95, 5048–5061 (2008).

28. Guo, Y., Martinez-Williams, C., Gilbert, K. A. & Rannels, D. E. Inhibition of gap junction communication in alveolar epithelial cells by 18α-glycyrrhetinic acid. Am J Physiol Lung Cell Mol Physiol 276, (1999).

29. Peppiatt, C. M. et al. 2-Aminoethoxydiphenyl borate (2-APB) antagonises inositol 1,4,5-trisphosphate-induced calcium release, inhibits calcium pumps and has a use-dependent and slowly reversible action on store-operated calcium entry channels. Cell Calcium 34, 97–108 (2003).

30. Ma, J. Block by ruthenium red of the ryanodine-activated calcium release channel of skeletal muscle. J Gen Physiol 102, 1031–1056 (1993).

31. Lizák, B. et al. Translocon pores in the endoplasmic reticulum are permeable to small anions. Am J Physiol Cell Physiol 291, (2006).

32. Corsi, A. K. & Schekmant, R. Mechanism of polypeptide translocation into the endoplasmic reticulum. J Biol Chem 271, 30299–30302 (1996).

33. Wulf, J. & Pohl, W. G. Calcium ion-flux across phosphatidylcholine membranes mediated by ionophore A23187. Biochim Biophys Acta 465, 471–485 (1977).

34. Karkempetzaki, A. I. & Ravid, K. Piezo1 and Its Function in Different Blood Cell Lineages. Cells 13, (2024).

35. Allbritton, N. L., Meyer, T. & Stryer, L. Range of messenger action of calcium ion and inositol 1,4,5-trisphosphate. Science 258, 1812–1815 (1992).

36. Moreno, A. P., Sáez, J. C., Fishman, G. I. & Spray, D. C. Human connexin43 gap junction channels. Regulation of unitary conductances by phosphorylation. Circ Res 74, 1050–1057 (1994).

37. Miroshnikova, Y. A. & Wickström, S. A. Mechanical Forces in Nuclear Organization. Cold Spring Harb Perspect Med 14, (2022).

38. Ferrera, D. et al. Lamin B1 overexpression increases nuclear rigidity in autosomal dominant leukodystrophy fibroblasts. FASEB J 28, 3906–3918 (2014).

39. Nava, M. M. et al. Heterochromatin-driven nuclear softening protects the genome against mechanical stress-induced damage. Cell 181, 1–18 (2020).

40. Damodaran, K. et al. Compressive force induces reversible chromatin condensation and cell geometry–dependent transcriptional response. Mol Biol Cell 29, 3039–3051 (2018).

41. Lomakin, A. J. et al. The nucleus acts as a ruler tailoring cell responses to spatial constraints. Science 370, (2020).

42. Enyedi, B., Jelcic, M. & Niethammer, P. The Cell Nucleus Serves as a Mechanotransducer of Tissue Damage-Induced Inflammation. Cell 165, 1160–1170 (2016).

43. Gong, M. C., Kinter, M. T., Somlyo, A. V. & Somlyo, A. P. Arachidonic acid and diacylglycerol release associated with inhibition of myosin light chain dephosphorylation in rabbit smooth muscle. J Physiol 486, 113 (1995).

44. Schanstra, J. P. et al. Systems biology identifies cytosolic PLA2 as a target in vascular calcification treatment. JCI Insight 4, (2019).

45. Wu, M., Wu, X. & De Camilli, P. Calcium oscillations-coupled conversion of actin travelling waves to standing oscillations. Proc Natl Acad Sci U S A 110, 1339–1344 (2013).

46. Shao, X., Li, Q., Mogilner, A., Bershadsky, A. D. & Shivashankar, G. V. Mechanical stimulation induces formin-dependent assembly of a perinuclear actin rim. Proc Natl Acad Sci U S A 112, E2595–E2601 (2015).

47. Khalilgharibi, N. et al. Stress relaxation in epithelial monolayers is controlled by the actomyosin cortex. Nat Phys 15, 839–847 (2019).

48. Mao, Y. & Wickström, S. A. Mechanical state transitions in the regulation of tissue form and function. Nature Reviews Molecular Cell Biology 2024 25:8 25, 654–670 (2024).

49. Casares, L. et al. Hydraulic fracture during epithelial stretching. Nat Mater 14, 343–351 (2015).

50. Colom, A. et al. A fluorescent membrane tension probe. Nat Chem 10, 1118–1125 (2018).

51. Huber, M. D. & Gerace, L. The size-wise nucleus: nuclear volume control in eukaryotes. J Cell Biol 179, 583–584 (2007).

52. Kim, J. K. et al. Nuclear lamin A/C harnesses the perinuclear apical actin cables to protect nuclear morphology. Nature Communications 2017 8:1 8, 1–13 (2017).

53. Tomba, C., Luchnikov, V., Barberi, L., Blanch-Mercader, C. & Roux, A. Epithelial cells adapt to curvature induction via transient active osmotic swelling. Dev Cell 57, 1257–1270.e5 (2022).

54. Jetta, D., Gottlieb, P. A., Verma, D., Sachs, F. & Hua, S. Z. Shear stress-induced nuclear shrinkage through activation of Piezo1 channels in epithelial cells. J Cell Sci 132, 1–8 (2019).

55. Phengchat, R. et al. Calcium ions function as a booster of chromosome condensation. Sci Rep 6, 1–10 (2016).

56. Aureille, J. et al. Nuclear envelope deformation controls cell cycle progression in response to mechanical force. EMBO Rep 20, e48084 (2019).

57. Elosegui-Artola, A. et al. Force Triggers YAP Nuclear Entry by Regulating Transport across Nuclear Pores. Cell 171, 1397–1410.e14 (2017).

58. Brun-Cosme-Bruny, M. et al. Microfluidic system for in vitro epithelial folding and calcium waves induction. STAR Protoc 3, 101683 (2022).

59. Long, J., Junkin, M., Wong, P. K., Hoying, J. & Deymier, P. Calcium wave propagation in networks of endothelial cells: model-based theoretical and experimental study. PLoS Comput Biol 8, (2012).

